# Type I-F CRISPR-Cas resistance against virulent phage infection triggers abortive infection and provides population-level immunity

**DOI:** 10.1101/679308

**Authors:** Bridget N.J. Watson, Reuben B. Vercoe, George P.C. Salmond, Edze R. Westra, Raymond H.J Staals, Peter C. Fineran

**Affiliations:** Department of Microbiology and Immunology, University of Otago, PO Box 56, Dunedin 9054, New Zealand; Department of Biochemistry, University of Cambridge, Cambridge CB2 1QW, United Kingdom; ESI, Biosciences, University of Exeter, Cornwall Campus, Penryn TR10 9FE, UK; Laboratory of Microbiology, Wageningen University and Research, 6708 WE Wageningen, The Netherlands; Bio-Protection Research Centre, University of Otago, Dunedin, New Zealand

## Abstract

Type I CRISPR-Cas systems are the most abundant and widespread adaptive immune systems of bacteria and can greatly enhance bacterial survival in the face of temperate phage infection. However, it is less clear how these systems protect against virulent phages. Here we experimentally show that type I CRISPR immunity of *Pectobacterium atrosepticum* leads to rapid suppression of two unrelated virulent phages, ΦTE and ΦM1. However, unlike the case where bacteria are infected with temperate phages, this is the result of an abortive infection-like phenotype, where infected cells do not survive the infection but instead become metabolically inactive and lose their membrane integrity. Our findings challenge the view of CRISPR-Cas as a system that protects the individual cell and supports growing evidence of an Abi-like function for some types of CRISPR-Cas systems.

---

To respond to the pressure of phage infection, bacteria have evolved various lines of defence^1-3^. The adaptive arm of these defences is provided by CRISPR-Cas, which provides immunity through CRISPR RNA guided cleavage of phage genomes^4,5^. CRISPR-Cas systems are incredibly diverse, and are currently classified into two major classes (1 and 2), six types (I to VI) and >30 subtypes^6,7^ (for recent reviews, see^4,5,8^). Crucially, recent studies revealed that at least some CRISPR-Cas variants, belonging to types VI and III, induce cell dormancy through collateral RNA cleavage following target recognition^9-13^. Furthermore, it is possible that type V systems induce cell death through ssDNA cleavage^14^. In contrast, the most abundant type I CRISPR systems, which make up around 60% of all CRISPR systems^15^, as well as the somewhat less common type II systems, immunity are thought to increase survival of infected individuals^16,17^. However, so far experimental studies on type I systems have almost exclusively focused on interactions between bacteria and filamentous phages or obligate killing mutants of temperate phages, and it is therefore less clear how bacteria with CRISPR immunity resist virulent phages. Here we explored this question using *Pectobacterium atrosepticum*, which carries a type I-F system, and two unrelated virulent phages as a model system. We found that CRISPR-Cas immunity reduced the number of cells that released phages and of those that produced progeny, the burst size was decreased. Infected cells did not survive phage infection, yet they reduced phage amplification, which provided protection at the population level. The observed CRISPR-Cas immunity phenotype to virulent phage infection has key implications for the way natural selection operates on these genes^18^ and is analogous to that observed for kin-selected altruistic defences such as abortive infection systems, which also provide population-level benefits at high individual cost.

## Results

### CRISPR-Cas reduces phage infectious centres and burst size

To investigate the outcomes of phage infection in the presence of CRISPR-Cas immunity, we examined the response to phage infection by *P. atrosepticum*, which contains a type I-F system^19,20^. We used two different phages, ΦTE and ΦM1, members of the *Myoviridae* and *Podoviridae*, respectively. Phage infectivity was assessed using strains with one or three spacers in the chromosomal CRISPR arrays, with or without phage-targeting spacers. CRISPR-Cas provided protection against ΦTE and ΦM1 infection, reducing the efficiency of plating (EOP) by at least 10-fold, with additional spacers increasing resistance to 10^5^-fold (Fig. 1A, Table S1). To determine what stage of phage reproduction was impeded, we investigated the effects of CRISPR-Cas on defined aspects of infection. CRISPR-Cas caused a decrease in the efficiency of centre of infection (ECOI) formation (Fig. 1B), meaning that for ΦTE, only 4 or 1% of infected cells released at least one infectious phage (for the 1× and 3×anti-ΦTE strains, respectively). Following ΦM1 infection, only 23 or 6% of cells released phages (for 1× and 3×anti-ΦM1 respectively). Next, one-step growth curves were performed to observe phage growth on the resistant hosts (Fig. S1 and Table S1). The average phage burst size was determined for each host and the number was significantly reduced by CRISPR-Cas (Fig. 1C). For ΦTE, both the 1× or 3×anti-ΦTE strains almost completely suppressed the burst and for ΦM1 it was reduced by >90% on the 3×anti-ΦM1 strain. As expected, adsorption was unaffected by CRISPR immunity (Fig. S1 and Table S1). Therefore, the *P. atrosepticum* type I-F CRISPR-Cas immunity reduced both the number of cells releasing phages and the average number of phages released per cell.

**Figure 1.**
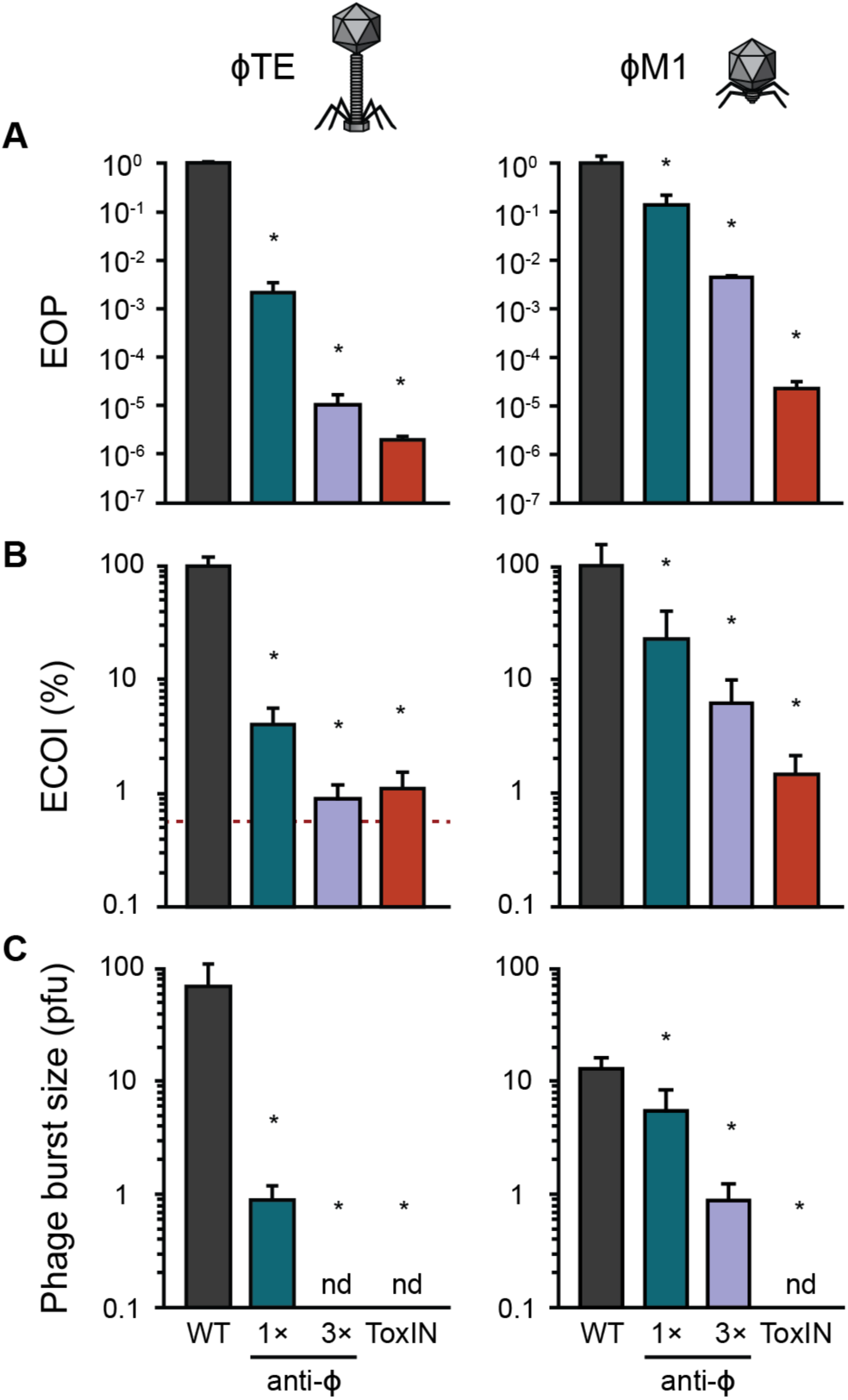
CRISPR-Cas reduces phage infectious centres and burst size. **A** Phage resistance (i.e. efficiency of plating (EOP)), **B** efficiency of centre of infection (ECOI) formation and **C** average burst size was assessed for the phage sensitive WT, cells with ToxIN and anti-Φ strains, with one (1×) or three (3×) spacers targeting ΦTE and ΦM1. Data shown is the mean +SD, nd = not detected. In **B** the red dashed line represents the background level of phages (in the other panels the limit of detection is below the axis limits). Statistical significance was calculated using one-way ANOVA using Dunnett’s multiple comparison test, comparing strains with targeting spacers to the control with no-targeting spacers. No significance was detected, unless indicated (* p ≤ 0.05). All data are provided in Table S1.

We previously characterised an Abi system in *P. atrosepticum*, ToxIN, which functions as a toxin-antitoxin system^21,22^. ToxIN provides protection against both ΦTE and ΦM1 phages, acting as an abortive infection mechanism, so we included ToxIN to compare the phenotypes provided by CRISPR-Cas and Abi immunity genes^21-23^. The ToxIN Abi system provided strong phage protection, reducing the EOP by 10^6^ and 10^5^-fold against ΦTE and ΦM1, respectively (Fig. 1A). For both phages, only 1% of phage infected cells harbouring ToxIN released any new viral progeny (Fig. 1B) and the average burst size was undetectable (Fig. 1C). As expected for a post-adsorption phage resistance mechanism, ToxIN had no effect on adsorption (Table S1). The outcomes of ToxIN and CRISPR-Cas-mediated immunity on the different aspects of infection were therefore qualitatively similar with respect to phage adsorption and amplification.

### The type I-F CRISPR-Cas system does not enable survival of infected cells

Next, we assessed cell survival of bacteria with CRISPR-Cas immunity upon infection with the virulent phages. Surprisingly, CRISPR-Cas immunity provided no enhancement in cell survival measured in viable count assays compared with the phage sensitive WT or the ToxIN Abi system (Fig. 2A), regardless of the multiplicities of infection (MOI) that were used (Fig. S2A). To further investigate cell survival, we assessed membrane integrity and cellular metabolic activity of phage infected cells (Fig. 2B and C, Fig. S2B and C). Phage infection led to significant reductions in both membrane integrity and cellular metabolism even in the presence of CRISPR-Cas or ToxIN immunity. As a control, surface mutants (i.e. bacteria carrying mutations in the phage receptor genes on the bacterial cell surface) were isolated that were resistant to either phage. As expected for adsorption inhibition, surface resistance against either phage resulted in cells retaining membrane integrity and metabolic activity upon phage challenge, but not when challenged with a phage that uses a different receptor (Fig. S3). Therefore, infected *P. atrosepticum* cells bearing type I-F CRISPR-Cas immunity limit phage propagation at the expense of the individual – akin to altruistic abortive infection.

**Figure 2.**
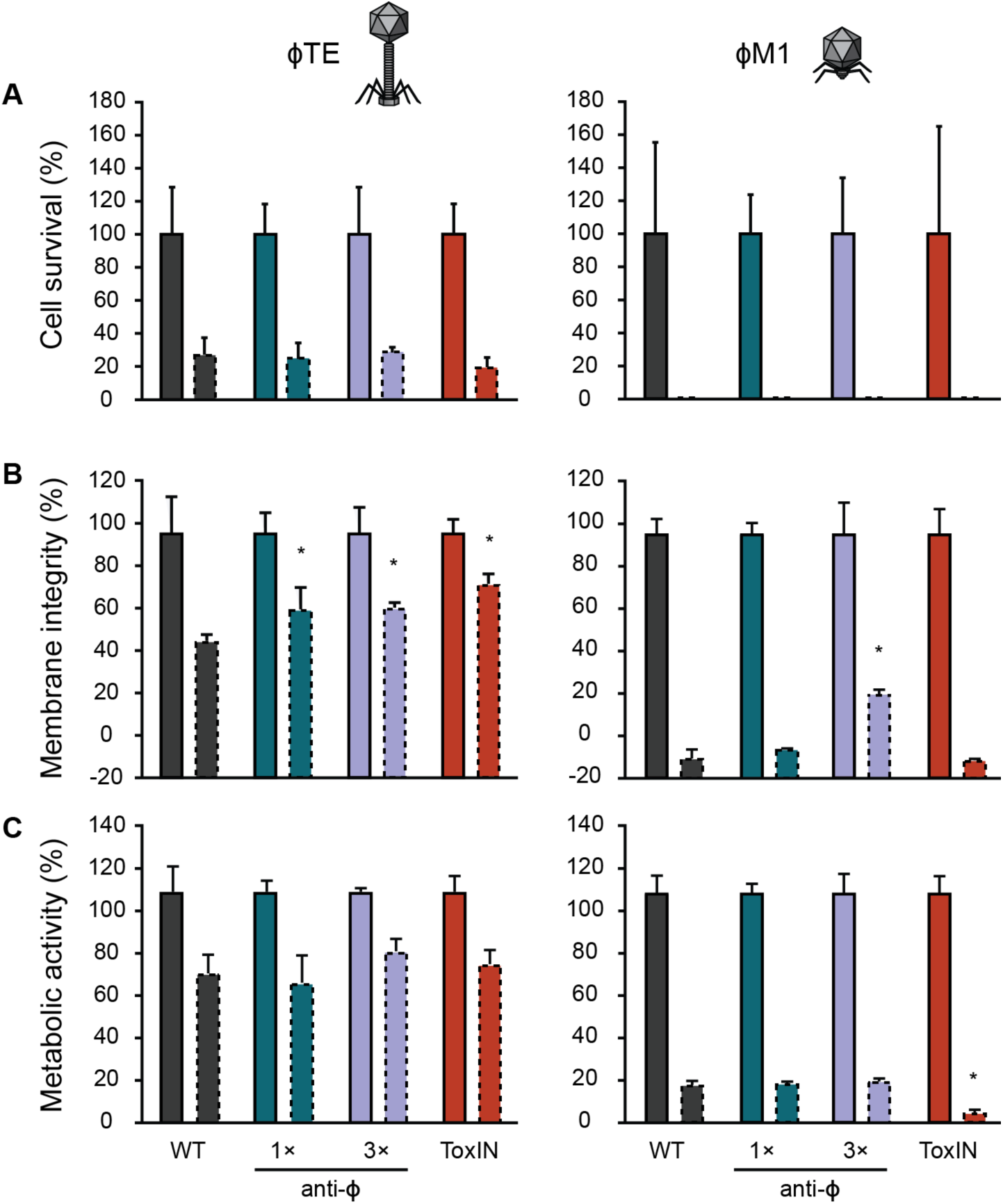
The type I-F CRISPR-Cas system does not enable survival of infected cells. **A** Cell survival was assessed for the WT, 1× and 3×anti-Φ strains, and ToxIN, using both ΦTE and ΦM1 (infected at an MOI of 2). **B** The percentage of cells with intact membranes was determined using LIVE/DEAD™ staining and **C** the percentage of metabolically active cells was assessed using the resazurin dye. For **B** and **C** cells were infected at an MOI of 2.5. Solid outline bars represent mock infected samples, dashed outline bars represent phage infected samples. Statistical significance was calculated using one-way ANOVA using Dunnett’s multiple comparison test, comparing strains with targeting spacers to the control with no-targeting spacers. No significance was detected, unless indicated (* p ≤ 0.05).

### Increased CRISPR-Cas resistance does not enhance survival of infected individuals

One possible explanation why CRISPR-Cas did not promote survival following infection could be due to an insufficient immune response, leading to incomplete phage clearance. Since the phage-targeting spacers are in CRISPR arrays carrying 30 (CRISPR1) and 11 (CRISPR2) other spacers, most effector complexes will be loaded with non-phage-targeting crRNAs. To explore if an increased abundance of Cas complexes loaded with phage-targeting crRNAs would result in survival of infected cells, phage targeting spacers were overexpressed from plasmids in the presence or absence of Cas overexpression (Fig. 3). Increased phage-targeting crRNAs significantly boosted phage resistance compared with chromosomal expression, and induction of Cas expression further enhanced resistance, by up to ∼10^4^-10^7^ fold compared to the WT (Fig. 3A). However, no marked restoration in cell survival was detected compared with the sensitive WT strain (Fig. 3B).

**Figure 3.**
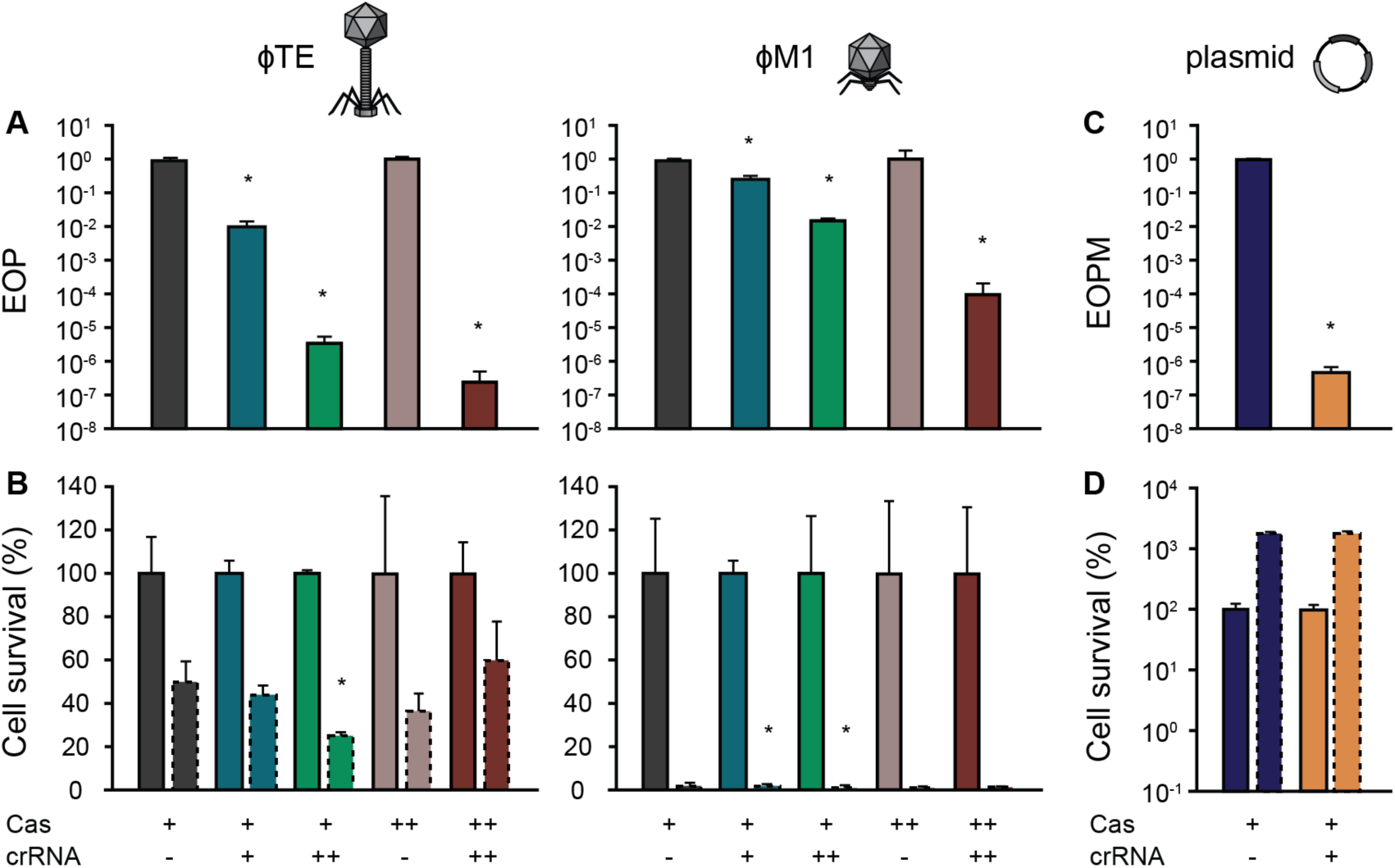
CRISPR-Cas overexpression increases resistance against phages but infected cells do not survive infection. **A** Phage resistance (EOP) and **B** cell survival was assessed for WT (with empty vector, pPF975) “Cas+ crRNA-”, 1×anti-Φ (PCF190 / PCF254 (with empty vector), chromosomally expressed) “Cas+ crRNA+”, 1×anti-Φ plasmid (WT carrying pPF1423 / pPF1421) “Cas+ crRNA++”, Cas overexpression (PCF610 (with empty vector)) “Cas++ crRNA-” and Cas overexpression with 1×anti-Φ plasmid (PCF610, pPF1423 / pPF1421) “Cas++ crRNA++”. Solid outline bars represent mock infected samples, dashed outline bars represent phage infected samples. **C** Efficiency of plasmid maintenance (EOPM) and **D** cell survival was assessed for strains carrying pTargeted (with the *expI* gene) and pControl (inducible mini CRISPR array with no anti-*expI* spacer) “Cas+ crRNA-” or pTargeted and pCRISPR (anti-*expI* spacer) “Cas+ crRNA+”. Solid outline bars represent CRISPR repressed samples, dashed outline bars represent CRISPR induced samples. Statistical significance was calculated using one-way ANOVA using Dunnett’s multiple comparison test, comparing strains with targeting spacers to the control with no-targeting spacers. The Cas overexpression and Cas overexpression with 1×anti-Φ plasmid strains, as well as the strains in **D**, were compared using an unpaired T-test. No significance was detected, unless indicated (* p ≤ 0.05).

While these data show that CRISPR-immune bacteria do not survive virulent phage infection even under artificially high CRISPR expression levels, it is unclear whether this is due to programmed cell death induced by CRISPR (analogous to that observed for type VI systems^10^, or due to the phage, which may express lethal genes prior to clearance of the infection. To explore this question, we examined the outcome of targeting plasmid DNA for the cells with CRISPR-Cas immunity (Fig. 3C and D). The *P. atrosepticum* CRISPR-Cas system effectively inhibits transformation and conjugation^24^, but those assays fail to assess the outcome for cells eliciting effective CRISPR immunity (they are killed by the antibiotic). To directly test whether plasmid targeting by the I-F system reduces cell survival in *P. atrosepticum*, we induced a mini-CRISPR array with a spacer targeting a plasmid and assessed total cell counts and plasmid loss. Plasmid targeting decreased cells bearing the plasmid by 10^6^-fold in 18 h but did not decrease total cell numbers. Hence, these experiments show that the Abi phenotype is phage-dependent, since cells survived plasmid targeting.

### CRISPR-Cas provides population-level protection at low phage doses

Even though an Abi-like phenotype is costly for the infected individual, they may be favoured by natural selection because of their population-level benefits if these are predominantly directed at clone mates (i.e. kin selection). To explore these kin-selected benefits, we compared population growth of cells carrying CRISPR-Cas or Abi under increasing phage pressures (increasing MOIs) (Fig. S4). Phage sensitive WT *P. atrosepticum* populations were susceptible to phages at any MOI. The phage effects on population growth were stronger and faster with increasing phage numbers, but even with an MOI of 0.0001, WT populations collapsed (Fig. 4A). As predicted for an Abi mechanism, cultures containing ToxIN grew with low phage doses, but when phages equalled or exceeded bacteria (MOI of 1 or higher) population growth was inhibited. Likewise, CRISPR-Cas immunity enabled population growth at low phage doses, but at higher MOIs, the populations either collapsed when infected with ΦM1, or became static when infected with ΦTE (Fig. 4A, Fig. S4). We predicted that CRISPR-Cas was providing population-level protection by reducing the phage epidemic. To test this, the effect of CRISPR-Cas on phage titres was determined (Fig. 4B). Both phages replicated extensively on the phage-sensitive WT bacteria, reaching ∼10^10^-10^11^ pfu ml^-1^ irrespective of the initial phage dosage (Fig. 4B). ToxIN reduced the population phage burden regardless of the initial phage abundance. CRISPR-Cas immunity limited the phage epidemic when initial viral abundance was low, but when initial phage numbers were higher, CRISPR was unable to suppress the phage burden. In summary, immunity provided by the type I-F CRISPR-Cas system enables population growth under low viral load by reducing virulent phage burden, but both CRISPR-Cas and ToxIN fail to cope with high phage pressures.

**Figure 4.**
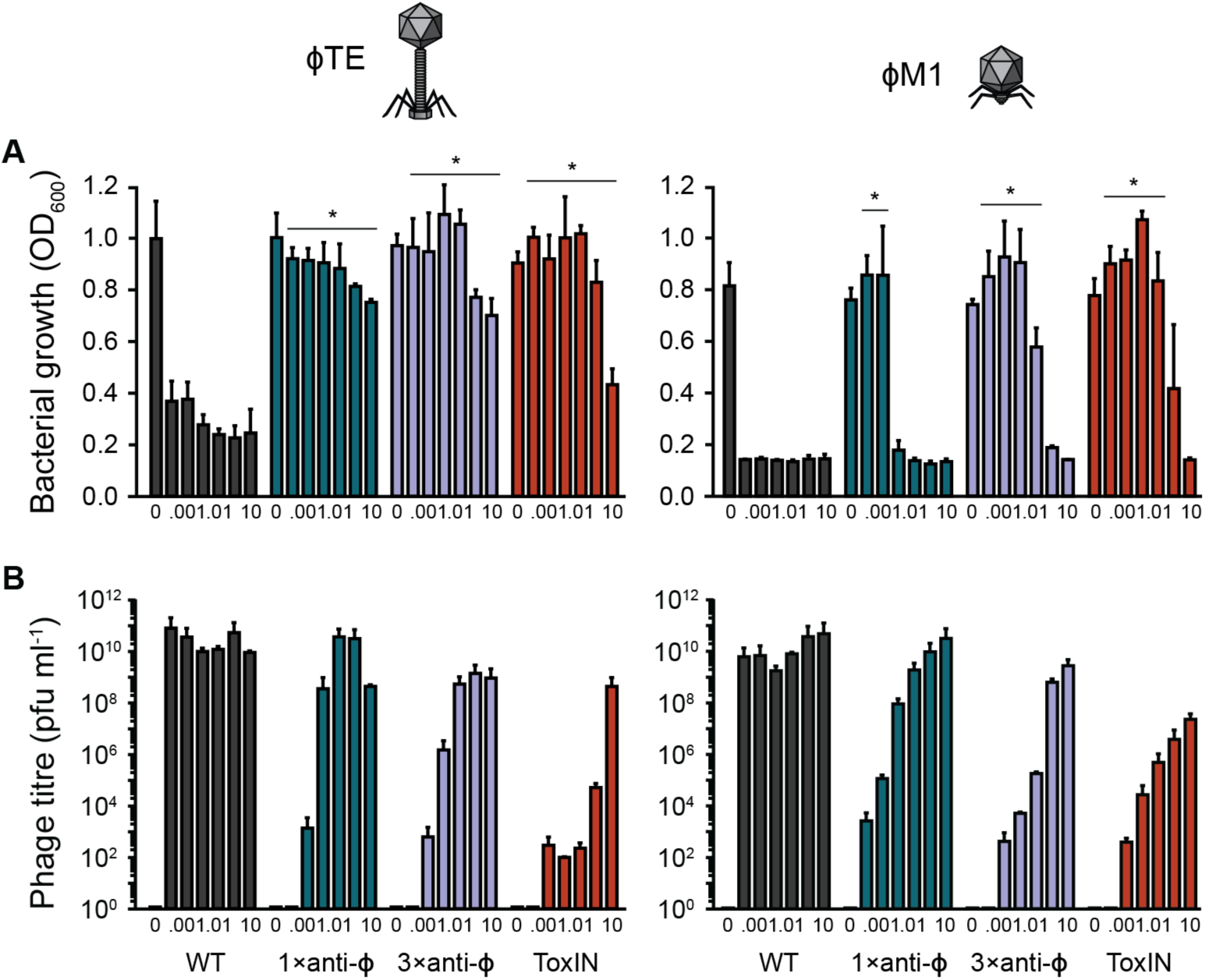
Populations of anti-Φ strains only grow at low phage doses. WT, 1× and 3×anti-Φ strains, and ToxIN were grown with phages, added at a range of doses (MOIs: 0 (buffer), 10^−^ 4, 10^-3^, 10^-2^, 10^-1^, 1, and 10). **A** The final bacterial growth levels (OD_600_) and **B** the final phage titres were determined after 16 h. Statistical significance was calculated using one-way ANOVA using Dunnett’s multiple comparison test, comparing strains with targeting spacers to the control with no-targeting spacers. No significance was detected, unless indicated (* p ≤ 0.05).

## Discussion

Here we show that the *P. atrosepticum* type I-F CRISPR-Cas system generates an immunity phenotype similar to that Abi systems, in which infected cells limit phage propagation and as a result, protect neighbouring cells from infection. CRISPR-Cas reduced phage infectivity, resulting in fewer infectious centres with reduced phage burst sizes and infected cells did not survive, became metabolically inactive and lost membrane integrity. However, population-level protection was achieved through CRISPR-Cas-mediated reduction in the phage epidemic. Although the CRISPR-Cas response to phage infection has been investigated in other systems, they are typically carried out with filamentous phages or virulent mutants of temperate phages, and thought to enhance survival of the infected individual^25-29^. Cell survival was demonstrated for *S. thermophilus* with a type II-A system^17^ and in other studies survival may be inferred since CRISPR adapted clones grow in the presence of phages^16,30-32^. The observation that the native *P. atrosepticum* system generates an Abi-phenotype upon infection with two virulent phages helps to explain previous observations that infection by T7 or T5 virulent phages targeted by the type I-E system of *Escherichia coli* slowed or inhibited bacterial growth^33^.

The ‘suicidal’ response of CRISPR-Cas to phage infection might occur through several mechanisms^34^. These include: activation of toxic domains found in some Cas proteins, such as Cas2^35^, off-target effects of the promiscuous RNA-targeting effector proteins from type III^11-13^ and type VI^10^ systems, and self-targeting due to increased spacer acquisition following CRISPR-Cas activation^36^. None of these models explain the Abi phenotype of the type I-F system in *P. atrosepticum*. For example, *P. atrosepticum* Cas2 does not have detectable nuclease (i.e. toxic) activity^37^ and although we have observed acquisition of self-targeting spacers, this is at a low frequency that is unlikely to significantly impact cell survival^38^. Indeed, PCR analysis of CRISPR array expansion following phage infection failed to detect spacer acquisition (data not shown). Instead, our data suggest that post-infection immunity by CRISPR-Cas response provides a window of time during which the virulent phage can express toxic phage products^33^. Temperate and filamentous phages can transmit both horizontally and vertically and therefore generally avoid immediate early expression of highly toxic genes as this would be associated with severe fitness trade-offs when the phage enters the lysogenic cycle. Early expressed virulent phage genes can lead to host DNA degradation, inhibition of host RNA polymerase and other effects^39,40^. Although the exact mechanism of host cell takeover for the two phages used in this study is unknown, ΦM1 encodes its own RNAP, suggesting a rapid host-takeover, and we have also shown that a gene responsible for triggering ToxIN immunity is toxic in *P. atrosepticum*^23^. Consistent with our phage-induced growth inhibition hypothesis, the Abi phenotype was absent during type I-F plasmid targeting.

In contrast to our findings, type I-E and I-F CRISPR systems can provide resistance against temperate and filamentous phages without apparent Abi phenotypes^16,41^. Chronic phage infection (M13) or obligately lytic temperate phage mutants (e.g. DMS3*vir*) do not rapidly, or strongly, manipulate bacterial physiology and therefore CRISPR immunity is sufficient to clear infection and protect the cell. Nonetheless, *P. aeruginosa* deployment of type I-F CRISPR-Cas causes a bacterial fitness cost, potentially due to decreased replication or repairing the damage following phage infection^16^. Despite this, CRISPR-Cas was advantageous to *P. aeruginosa* through the generation of diverse immunity against phages^42^. It is not clear how cells containing a CRISPR system that functions through an Abi-like phenotype can acquire new spacers. However, by analogy to work in a type II system, the type I-F system may acquire spacers during infection by defective phages^28^, which might enable phage-resistance to arise when bacteria are growing in a structured environment^18^. Indeed, Abi systems evolve in spatially structured environments where clone mates benefit directly^18,43,44^ and we predict that the *P. atrosepticum* CRISPR-Cas system will be beneficial under such conditions.

The Abi phenotype of this type I-F system strengthens the view that CRISPR-Cas immunity can sometimes coming at the expense of the individual, but providing a benefit for the population^10,45,46^. The nature of the invading element, the relative efficiency of resistance and type of CRISPR-Cas system are likely to influence whether CRISPR-Cas provides protection to the infected individual and the population, or just to the population via Abi. We predict that virulent phages are more likely to elicit Abi phenotypes, whereas temperate phages or other mobile elements will be more likely be cleared and result in cell survival. These outcomes likely fall on a spectrum determined by the invader vs host immune strength and will need to be factored in to ecological and evolutionary analyses of CRISPR-Cas immunity.

## Materials and Methods

### Bacterial strains, plasmids, and culture conditions

Bacterial strains and plasmids used in this study are given in Table S2. *P. atrosepticum* SCRI1043^47^ was grown at 25°C and *Echerichia coli* at 37°C in Lysogeny Broth (LB) at 180 rpm or on LB-agar (LBA) plates containing 1.5% (w v^-1^) agar. Minimal media contained 40 mM K_2_HPO_4_, 14.6 mM KH_2_PO_4_, 0.4 mM MgSO_4_, 7.6 mM (NH_4_)_2_SO_4_ and 0.2% (w v^−1^) glycerol. When required, media were supplemented with ampicillin (Ap, 100 µg ml^− 1^), kanamycin (Km; 50 µg ml^− 1^), isopropyl-ß-D-thiogalactopyranoside (IPTG, 0.1 mM), glucose (glu, 0.2% (v v^− 1^)) and arabinose (ara, 0.2% (v v^− 1^)). Bacterial growth was measured in a Jenway 6300 spectrophotometer at 600 nm (OD_600_). All experiments were performed in a minimum of biological triplicates and data shown is the mean + standard deviation.

### Phage storage and titration

The phages, ΦTE^22^ (genome size of ∼142 kb) and ΦM1^23,48^ (genome size of ∼43 kb), were stored in phage buffer (10 mM Tris-HCl pH 7.4, 10 mM MgSO_4_ and 0.01% w v^-1^ gelatin). Phage stocks were titrated by serially-diluting phages in phage buffer, adding to 100 µl of *P. atrosepticum* culture (pre-grown in 5 ml LB overnight) in 4 ml top LBA (0.35% (ΦTE) and 0.5% (ΦM1) agar) and pouring onto LBA plates. Plates were incubated at 25°C overnight, plaques were counted and the titre determined as plaque forming units (pfu) ml^− 1^. Efficiency of plating (EOP) was calculated as: (pfu ml^− 1^ (test strain) / pfu ml^− 1^ (control strain, *P. atrosepticum*)). For the following assays (excluding the assays with the crRNA and Cas overexpression), strains carried the vector, pBR322, to control for ToxIN (which is on the pBR322 derivative, pTA46).

### Efficiency of centre of infection assays (ECOI)

Overnight cultures were OD-adjusted and 1 ml was used to inoculate a 25 ml culture in a 250 ml flask, for a starting OD_600_ of 0.1. Cells were grown until early stationary phase (OD_600_ of ∼0.3) before 10^9^ total phages (∼ 4 × 10^7^ pfu ml^− 1^) were added at a multiplicity of infection (MOI) of ∼0.1 and cultures were incubated with shaking for 20 min. Aliquots of 1 ml were extracted, washed twice in 1×phosphate-buffered saline (PBS), diluted and plated in top LBA with *P. atrosepticum* before the infected cells starting lysing. The pfu ml^− 1^ was determined for each strain and since each plaque was formed from the phages released from an individual cell, the titre represents the number of infectious centres formed. The ECOI was calculated as (pfu ml^− 1^ (test strain) / pfu ml^− 1^ (control strain, *P. atrosepticum*)). Spontaneous Φ-resistant surface mutants, PCF333 and PCF334, were included to control for unadsorbed phages.

### One-step growth curves

Overnight cultures were OD-adjusted and 1 ml was used to inoculate a 25 ml culture in a 250 ml flask, for a starting OD_600_ of 0.1. Cells were grown until early exponential phase (OD_600_ of 0.25-0.35) and 10^9^ total phages (∼ 4 × 10^7^ pfu ml^− 1^) were added, for an MOI of ∼0.1. Duplicate samples were taken at various timepoints, until 70 min post infection. One sample was plated immediately (non-treated sample, free phages and phage-infected cells), while the second was added to phage buffer containing chloroform (treated sample, free phages and phage accumulated inside infected cells), which lysed the cells, allowing the assessment of the total number of mature phages at each time point. Samples were diluted in phage buffer and plated in top LBA with *P. atrosepticum*. Phage adsorption over time was determined from the treated samples using the equation ((pfu ml^− 1^ (t=0) - pfu ml^− 1^ (t=0 to 70) / pfu ml^− 1^ (t=0)). The average phage burst size was also calculated from the treated samples, as number of phages released ((pfu ml^− 1^ (t=70) - pfu ml^− 1^ (t=30)) / the number of cells infected ((pfu ml^− 1^ (t=0) - pfu ml^− 1^ (t=30)). The latent period was determined from the treated samples as was defined as the time before the phage burst starts.

### Cell survival assays

Cells were grown to OD_600_ ∼0.3 and for each culture, 1 ml was transferred into two universals. One culture was infected with phages at a MOI of ∼2, while the other was mock infected, with phage buffer. Cultures were shaken at 180 rpm for 20 min for phages to adsorb and then cells were pelleted and resuspended in PBS to remove unadsorbed phages. Finally, cells were diluted and 100 µl samples were plated prior to the phage burst (40 min). Cell survival was calculated as (colony forming units (cfu) ml^− 1^ (phage treated sample) / cfu ml^− 1^ (mock treated sample).

To assess cell survival at a range of MOIs, 100 µl of each exponential phase culture was aliquoted into eight wells of a 96-well flat-bottomed plate for the addition of 10 µl phages at seven MOIs as well as a mock infection control (phage buffer). Cultures were shaken for 20 min for phages to adsorb, and to reduce the burden of secondary infection, a viricidal solution called TEAF (per ml: 680 µl of 4.3 mM FeS0_4,_ 320 µl 7.5% (w v^-1^) green tea solution (filter-sterilised)^49^) was then added, at a ratio of 75% (v v^− 1^) to each culture. The cultures were then diluted, more TEAF was added to each dilution and cells were plated as 5 µl spots. Survival for the ΦTE-infected cells was higher than predicted from the MOIs used, suggesting that despite high adsorption rates (Table S1), the phage was not able to infect well in these assays with high phage doses.

### LIVE/DEAD staining for membrane activity

Cell membrane integrity was assessed using the LIVE/DEAD™ *Bac*Light™ bacterial viability kit, consisting of two nucleic acid stains, syto-9 and propidium iodide (Life technologies^(tm)^). Cultures were prepared for live/dead staining as described above for the cell survival assays performed at a range of MOIs. Cells were infected for one hour, to allow for one complete round of infection, before being stained, according to the manufacturers’ instructions. Culture fluorescence was measured using a Thermo Scientific™ Varioskan™ plate reader, with excitation / emission wavelengths of 485 / 530 nm for styo-9 and 485 / 630 nm for propidium iodide. Cultures of exponentially growing cells and cells killed with 70% isopropanol were combined at different ratios to generate a standard curve, from which the percentage of cells with intact membranes at each phage MOI could be determined.

### Resazurin assays for cell activity

For assays assessing cell activity after one round of phage infection, cultures were prepared as described above for the cell survival assays performed at a range of MOIs. Cells were infected for one hour before resazurin solution was added at a final concentration of 0.005% (w v^− 1^). Cellular oxidoreductases reduce the blue indicator to resorufin, which is pink. Resorufin fluorescence was measured 30 min after it was added using a Thermo Scientific™ Varioskan™ plate reader with excitation / emission wavelengths of 510 / 535 nm. Cells for the standard curve were prepared as described for the live/dead staining, from which the percentage of metabolically active cells at each MOI was determined. Cell activity was assessed, following the 16 h growth assays, in the same way.

### Isolation of spontaneous phage-resistant surface mutant strains

ΦTE and ΦM1 were plated on *P. atrosepticum* and cells from colonies that formed in the centre of plaques were streaked to single colonies. Since ΦTE is flagella-trophic^50^, clones isolated from plates with ΦTE were patched onto tryptic swimming agar (10 g Bacto tryptone, 5 g NaCl, 3 g agar, per litre) to assess flagella-mediated swimming. A clone that did not swim (PCF333, Table S2) was resistant to ΦTE, but sensitive to ΦM1, which does not use the flagella for infection, suggesting that it was a surface mutant. A clone isolated from a ΦM1 plaque (PCF334, Table S2) was ΦM1-resistant, but sensitive to ΦTE.

### Construction of the plasmids expressing crRNAs

Spacers present in strains targeting ΦTE (PCF190) and ΦM1 (PCF254) were cloned into pPF975. Overlapping primers containing the spacer sequences were annealed and ligated into the BsaI site in the mini CRISPR array (repeat-repeat loci) as previously described^51^ to form the plasmids, pPF1421 and pPF1423 (Table S2). Oligonucleotide sequences are listed in Table S3. All plasmids used in this study were confirmed by sequencing.

### Construction of the cas overexpression strains

The chromosomal *cas* overexpression strain (PCF610) was made by conjugating the suicide vector, pPF1814, into *P. atrosepticum.* The vector, pPF1814 was constructed as follows: pSEVA511 was digested with NotI and ligated with the T5/*lac* promoter and multiple cloning site (MCS) from pQE-80L-stuffer, which had been amplified with PF3494 and PF3495 and digested with NotI. The *lacI* gene was amplified from pQE-80L-stuffer (PF2511, PF2512) and ligated into the MCS at XmaI and SalI sites. Finally, the first 500 bp of *cas1* was amplified using PF357 and PF669 and ligated into EcoRI and XmaI sites in the MCS.

### Plasmid targeting assay

The effect of plasmid targeting on cell survival was assessed using a two-plasmid setup. The first plasmid was either a control vector (pControl, pPF445, Ap^R^) with an inducible mini CRISPR array with a single repeat or pCRISPR (pPF452, Ap^R^) carrying a spacer targeting *expI*. The second plasmid was pTargeted (pPF459, Km^R^), which carried the targeted *expI* gene. pTargeted was made by PCR-amplifying *expI* from *P. atrosepticum* with PF314 and PF317, digesting the product with BamHI and PstI and ligating the product into the same sites in pPF260 (Km^R^-pQE-80L derivative). pControl and pCRISPR were made previously^52^. *P. atrosepticum* Δ*expI* (PCF81) was co-transformed with pTargeted and pCRISPR, or pControl, under CRISPR repressing conditions (0.2% glu) with both antibiotics (Km and Ap). These strains were for 6 hours in LB, 0.2% glu, Ap + Km with shaking. Cells were pelleted by centrifugation, washed and the culture was split into two samples, repressed (0.2% glu and Ap) and induced CRISPR conditions (0.2% ara and Ap). Following growth for a further 18 h, cells were plated onto Ap (for total cell counts) and Km (for targeted vector-containing cell counts). Efficiency of plasmid maintenance was calculated from the Km counts as (cfu ml^− 1^ (pCRISPR) / cfu ml^− 1^ (pControl)). Cell survival was calculated for each strain as (cfu ml^− 1^ (induced) / cfu ml^− 1^ (repressed). The cell counts for the induced CRISPR conditions were higher because the growth rate of *P. atrosepticum* was increased with supplemented arabinose.

### Bacterial population growth assays

*P. atrosepticum* cultures were grown to an OD_600_ of 0.3 and 100 μl was transferred to each well (of a 96-well plate). Phages were added in 10 μl at multiplicities of infection (MOIs) ranging from 0.0001 to 10 and cultures were grown in a Thermo Scientific™ Varioskan™ plate reader with shaking at 480 rpm. Cell density was monitored for 16 h, measuring OD_600_ every 12 min. Following growth, final phage titres were determined by chloroform treating the bacterial cultures and titrating the phages. The data were processed using GraphPad Prism to generate restricted cubic spline curves (324 points were calculated).

### Data availability statement

The data that support the findings of this study are available from the corresponding author upon reasonable request.

## Acknowledgements

This work was supported by a Rutherford Discovery Fellowship from the Royal Society of New Zealand (PCF), the Marsden Fund, RSNZ, the Bio-protection Research Centre (Tertiary Education Commission), a University of Otago Doctoral Scholarship (to BNJW), a Veni grant from the Netherlands Organization for Scientific Research (NWO) [016.Veni.171.047 to RHJS] and a Health Sciences Career Development Award from the University of Otago (to RHJS). The funders had no role in study design, data collection and interpretation, or the decision to submit the work for publication. Thanks to Josh Ramsay for providing plasmid pQE-80L-stuffer. We thank members of the Fineran laboratory for useful discussions.

