## Supplementary Information for "Type I-F CRISPR-Cas resistance against virulent phage infection triggers abortive infection and provides population-level immunity"

### Supplementary Figures

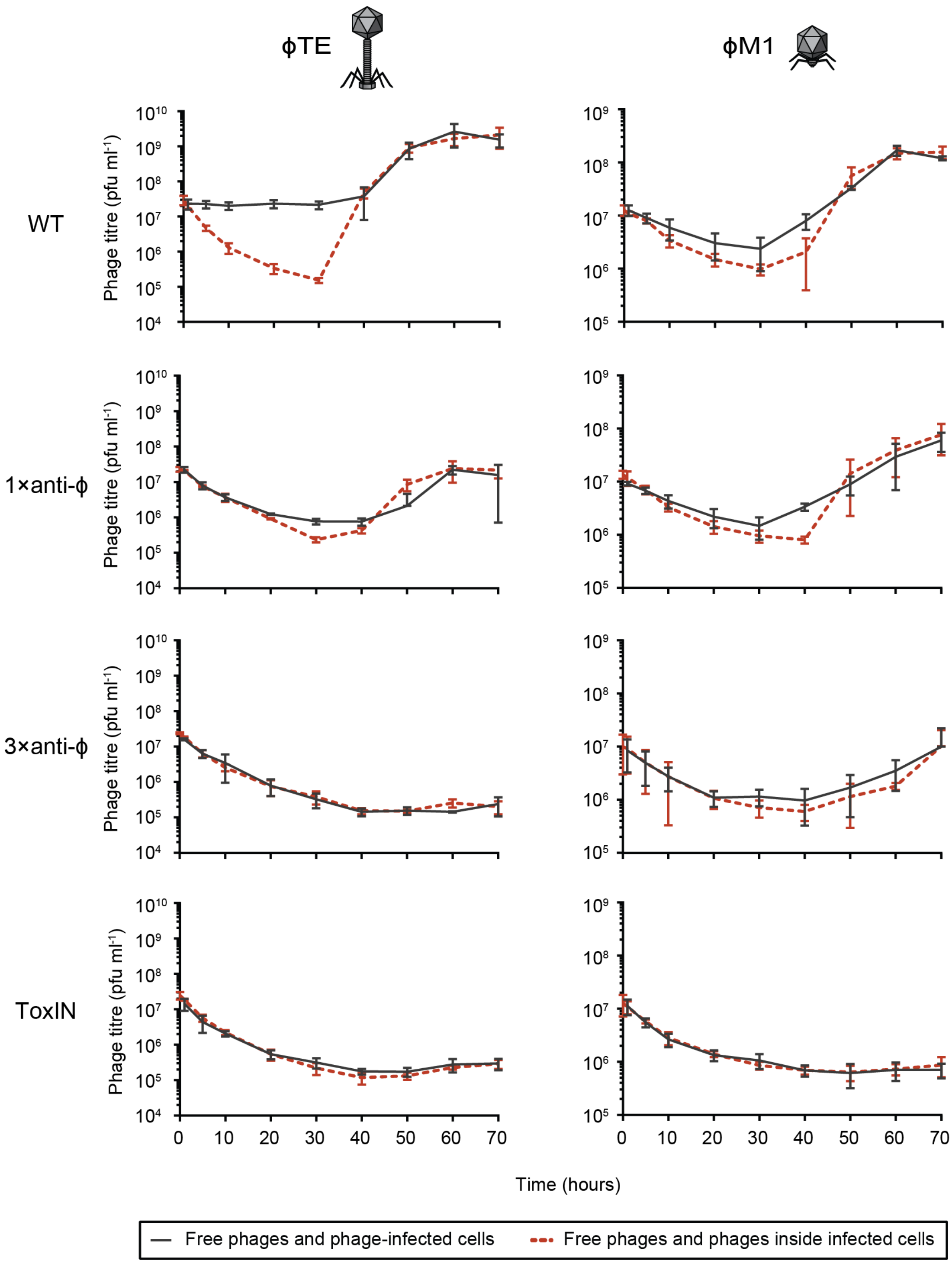


**Figure S1. One-step growth curves provide insight into adsorption and phage burst size.** Assays were performed on strains infected at an MOI of ~0.1. Samples were non-treated (free phages and phage-infected cells, black line) or treated with chloroform (free phages and phages accumulated inside infected cells, red dashed line). Phage burst size and adsorption data was calculated for Fig. 1 and Table S1.

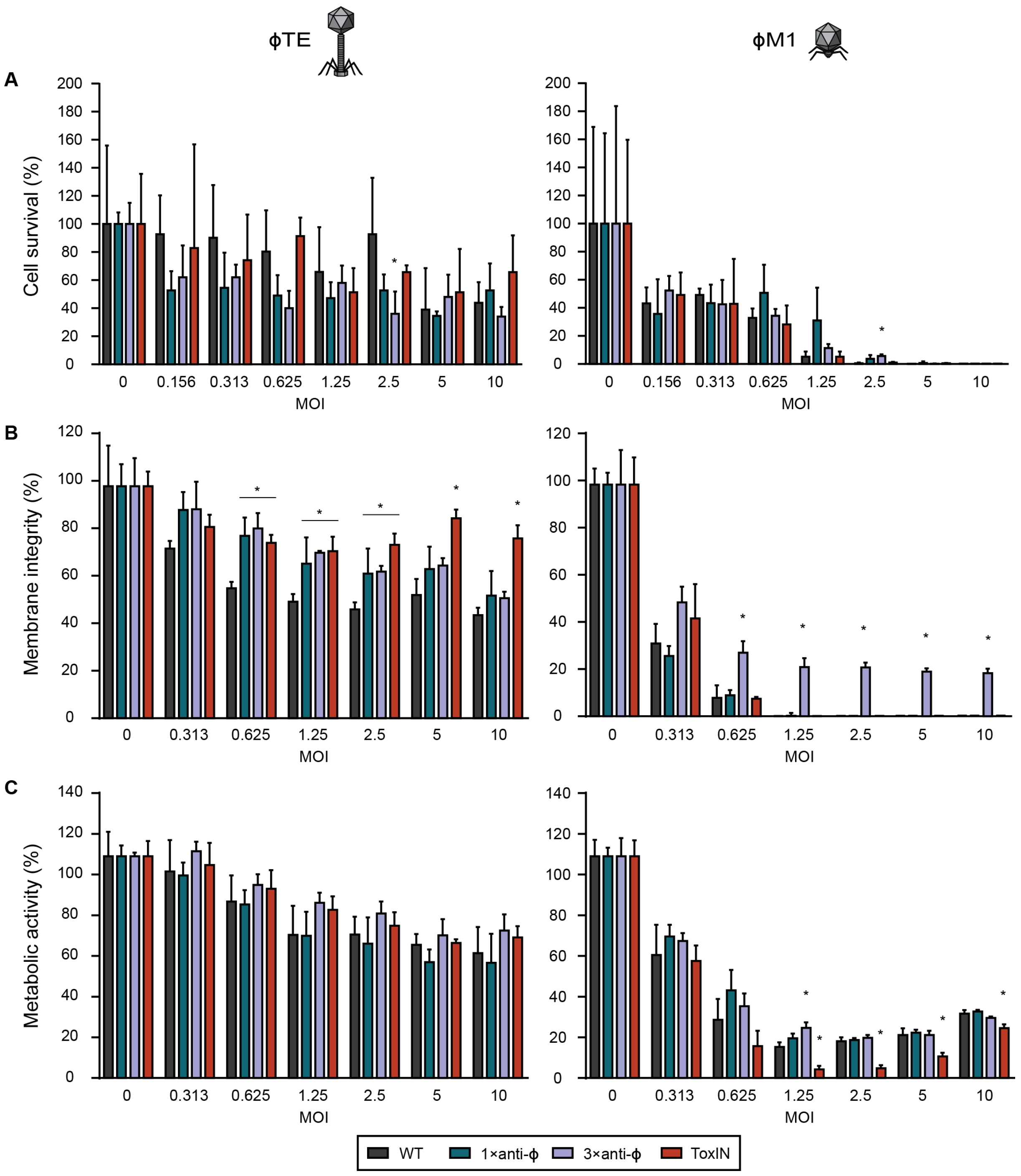


**Figure S2. CRISPR-Cas immunity does not promote cell survivial at a range of MOIs. A** Cell survival, **B** membrane integrity and **C** metabolic activity was assessed at a range of MOIs for WT, 1× and 3×anti-ϕ strains and ToxIN, using both ϕTE and ϕM1. Statistical significance was calculated using one-way ANOVA using Dunnett's multiple comparison test, comparing strains with targeting spacers to the control with no-targeting spacers. No significance was detected, unless indicated (* p ≤ 0.05).

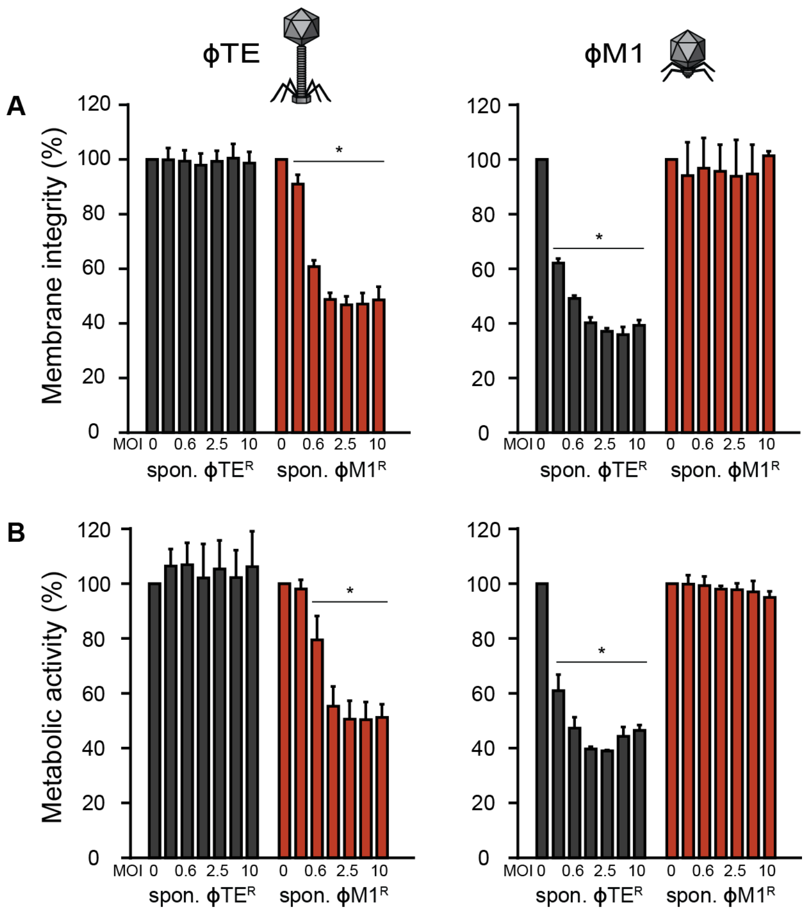


**Figure S3 Spontaneous phage-resistant mutants are active in the presence of phages.** Spontaneous ϕTE$^R$ and ϕM1$^R$ mutants were infected with ϕTE and ϕM1 at different MOIs (0, 0.3, 0.6, 1.25, 2.5, 5 and 10) and **A** membrane integrity **B** cell activity levels was assessed following one round of infection. Statistical significance was calculated using one-way ANOVA using Dunnett's multiple comparison test, comparing the phage-infected samples to the uninfected sample for each strain. No significance was detected, unless indicated (* $p \leq 0.05$).

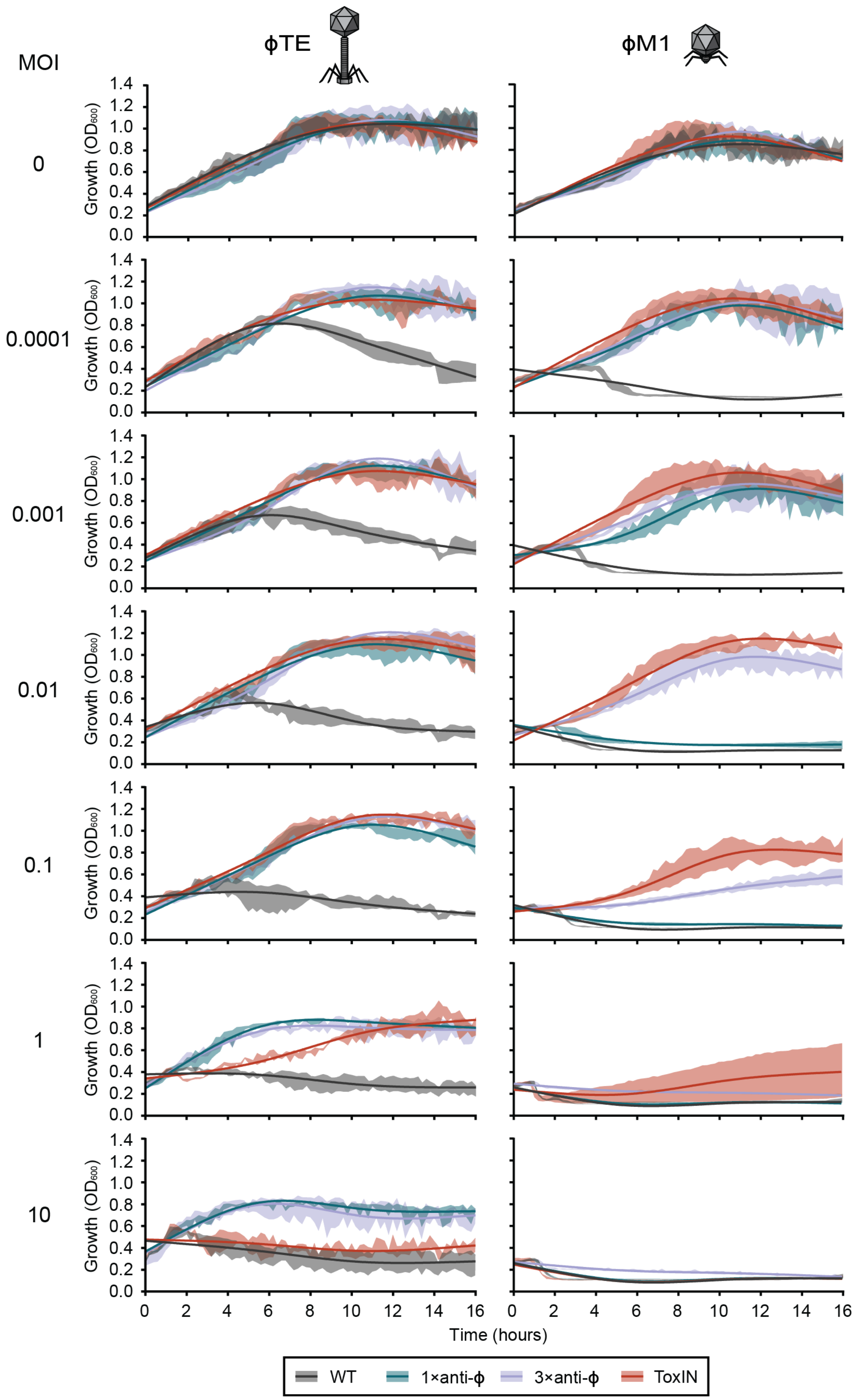


**Figure S4 Anti-ϕ strains grow in the presence of phages up to a MOI of 1.** Strains were grown in the presence of phages at different MOIs and $OD_{600}$ measurements were taken every 12 min for 16 hours. Solid lines: restricted cubic spline curve of the $OD_{600}$ values, shaded colour: one SD of the mean $OD_{600}$. These are the full data from what is presented in Fig. 4.

### Supplementary Tables

**Table S1.** Characteristics of phages ϕTE and ϕM1.

| Phage/ host | EOP | ECOI (%) | Latent period (min) | Adsorption (%) | Burst size (phages) |
|---|---|---|---|---|---|
| **ϕTE** | | | | | |
| WT | $1.0\times10^{0} \pm 6.0\times10^{-2}$ | 100 ±18.7 | 30 ±0 | 99 ±0.00 | 75 ±44 |
| 1×anti-ϕTE | $2.0\times10^{-3} \pm 1.2\times10^{-3}$ | 4.1 ±1.6 | 33 ±6 | 99 ±0.00 | 1 ±0 |
| 3×anti-ϕTE | $1.1\times10^{-5} \pm 5.6\times10^{-6}$ | 0.9 ±0.3 | n/d | 98 ±0.01 | <1 |
| ToxIN | $1.9\times10^{-6} \pm 4.1\times10^{-7}$ | 1.1 ±0.4 | n/d | 99 ±0.00 | <1 |
| **ϕM1** | | | | | |
| WT | $1.0\times10^{0} \pm 4.2\times10^{-1}$ | 100 ±50.7 | 37 ±6 | 92 ±0.04 | 13 ±3 |
| 1×anti-ϕM1 | $1.5\times10^{-1} \pm 6.5\times10^{-2}$ | 22.5 ±17.4 | 37 ±6 | 93 ±0.03 | 6 ±3 |
| 3×anti-ϕM1 | $4.7\times10^{-3} \pm 2.1\times10^{-4}$ | 6.3 ±3.5 | 40 ±0 | 90 ±0.08 | 1 ±0 |
| ToxIN | $2.3\times10^{-5} \pm 7.4\times10^{-6}$ | 1.5 ±0.6 | n/d | 93 ±0.02 | <1 |

Data shown is the mean ±SD. n/a not applicable. n/d no data the pfu values continue to decrease and there was no detectable phage burst.

**Table S2.** Bacterial strains and plasmids used in this study.

| Strain/Plasmid | Relevant Genotype/Phenotype | Reference |
|---|---|---|
| **Strains** | | |
| ***Escherichia coli*** | | |
| DH5α | F-, φ80Δ*dlacZ*M15, Δ(*lacZYA–argF*)U169, *endA1*, *recA1*, *hsd*R17 ($r_K^- m_K^+$), *deoR*, *thi-1*, *supE*44, λ-, *gyr*A96, *rel*A1 | Gibco/BRL |
| ST18 | *recA, pro, hsdR, recA*::RP4-2-Tc::Mu, λ*pir*, Tmp$^R$, Sp$^R$, Sm$^R$, Δ*hemA* | [1] |
| ***Pectobacterium atrosepticum*** | | |
| SCRI1043 | Wild type (WT) | [2] |
| PCF81 | SCRI1043 Δ*expI*::*cat*, Cm$^R$ | [3] |
| PCF188 | SCRI1043 with 3x anti-ϕTE spacers (in CRISPR1+2) | [4] |
| PCF190 | SCRI1043 with 1x anti-ϕTE spacer (in CRISPR1) | [5] |
| PCF254 | SCRI1043 with 1x anti-ϕM1 spacer (in CRISPR1) | [5] |
| PCF256 | SCRI1043 with 3x anti-ϕM1 spacers (in CRISPR1+2) | [5] |
| PCF333 | SCRI1043 with spontaneous ϕTE$^R$ | This study |
| PCF334 | SCRI1043 with spontaneous ϕM1$^R$ | This study |
| PCF610 | SCRI1043 with integrated pPF1814 for *cas* operon overexpression | This study |
| **Plasmids** | | |
| pBR322 | Cloning vector, ColE1 ori, Tc$^R$, Ap$^R$ | [6] |
| pPF260 | pQE-80L derivative with RP4 oriT, Km$^R$ | [7] |
| pPF445 (“pControl”) | mini-CRISPR with 1 repeat, pBAD30-derivative (aka pC1-16), p15a ori, Ap$^R$ | [3] |
| pPF452 (“pCRISPR”) | mini-CRISPR with single spacer targeting *expI*, pPF445 -derivative (aka pE1-16)), Ap$^R$ | [3] |
| pPF459 (“pTargeted”) | pPF260-derivative with *P. atrosepticum expI* gene, Km$^R$ | This study |
| pPF975 | pPF260-derivative, IPTG-inducible CRISPR locus for expressing crRNAs, Km$^R$ | [8] |
| pPF1421 | pPF975-derivative with the spacer from PCF254 | This study |
| pPF1423 | pPF975-derivative with the spacer from PCF190 | This study |
| pPF1814 | pSEVA511-derivative with T5/*lac* promoter, MCS and *lacI* from pQE-80L-stuffer, and 500 bp of *cas1* | This study |
| pQE-80L-stuffer | pQE-80L (Qiagen) with the 6His removed by digestion with EcoRI and BamHI and these sites restored, Ap$^R$ | Josh Ramsay; unpublished |
| pSEVA511 | R6K ori, Tc$^R$ | [9] |
| pTA46 | pBR322-derivative containing *toxIN*, Ap$^R$ | [10] |

**Table S3.** Oligonucleotide sequences used in this study

| Name | Sequence (5'-3') | Description |
|---|---|---|
| PF210 | GTCATTACTGGATCTATCAACAGG | R 100 bp downstream of CRISPR locus in pPF975 |
| PF314 | TTTGGTACCGGATCCGTGGCAATGATTACTCCATC | F for amplifying *expI* from *P. atrospeticum* (BamHI) |
| PF317 | TTTTCTAGACTGATGAATGGGTGAATCTC | R for amplifying *expI* from *P. atrospeticum* (XbaI) |
| PF357 | GACGAATTCTTACGGAAGAAAATACATTATGG | F for amplifying *cas1* N-terminal (EcoRI) |
| PF669 | TTTCCCGGGAAAGGTAAAGCGCGATTCAC | R for amplifying 500 bp into *cas1* (XmaI) |
| PF2511 | TCTCCCGGGAGGCATCAAATAAAACGA | F for amplifying *lacI* from pQE-80L (XmaI) |
| PF2512 | TCTGTCGACACACCATCGAATGGTGCA | R for amplifying *lacI* from pQE-80L (SalI) |
| PF2565 | GAAAACTAGCGTCTGTAGTGGGTCGTTGTGCAAGTAG | F for cloning PCF254 spacer into pPF975 |
| PF2566 | TGAACTACTTGCACAACGACCCACTACAGACGCTAGT | R for cloning PCF254 spacer into pPF975 |
| PF2569 | GAAATGACACAGCCAACGCCCTGAAAATCGGCACAGG | F for cloning PCF190 spacer into pPF975 |
| PF2570 | TGAACCTGTGCCGATTTTCAGGGCGTTGGCTGTGTCA | R for cloning PCF190 spacer into pPF975 |
| PF3494 | TTTGCGGCCGCTCGTCTTCACCTCGAGAAATC | F for amplifying pQE-80L MCS (NotI) |
| PF3495 | TTTGCGGCCGCGTCATTACTGGATCTATCAACAGG | R for amplifying pQE-80L MCS (NotI) |
